# Pneumococcal vaccine impacts on the population genomics of non-typeable *Haemophilus influenzae*

**DOI:** 10.1101/228791

**Authors:** David W. Cleary, Vanessa T. Devine, Denise E. Morris, Karen L. Osman, Rebecca A. Gladstone, Stephen D. Bentley, Saul N. Faust, Stuart C. Clarke

## Abstract

Between 2008/09 and 2012/13 the molecular epidemiology of non-typeable *Haemophilus influenzae* (NTHi) carriage in children <5 years of age was determined; a period that included pneumococcal conjugate vaccine (PCV) 13 introduction. Significantly increased carriage in post-PCV13 years was observed and lineage-specific associations with *S. pneumoniae* were observed before and after PCV13 introduction. NTHi were characterised into eleven discrete, temporally stable lineages, congruent with current knowledge regarding the clonality of NTHi. This increase could not be linked to the expansion of a particular clone and demonstrates different dynamics to before PCV13 implementation during which time NTHi co-carried with vaccine serotype pneumococci.

## Introduction

Non-typeable *Heamophilus influenzae* (NTHi) is a Gram-negative, human nasopharyngeal bacterium. Most research has leant towards the capsulated *Haemophilus influenzae,* in particular to serotype B, due to its capacity for causing severe invasive infections such as meningitis and septicemia^1^. *H. influenzae* type b (Hib) clinical disease has reduced with the widespread use of specific conjugate vaccines^1^. The global mortality and morbidity from NTHi, which predominantly replaced serotype B in nasopharyngeal carriage^2^ is significant. This includes the association with acute otitis media (AOM)^3^ and in exacerbations of chronic lung conditions such cystic fibrosis^4,5^ and chronic obstructive pulmonary disease^6^. Importantly NTHi is now also the leading cause of invasive *H. influenzae* disease. Examining the burden in 12 EU/EEA countries between 2007 and 2014, NTHi accounted for 78% of the 8,781 cases of invasive disease with the burden highest in infants and those ≥60 years of age^7^. The former increase was largely down to a concerning 6.2% (95% CI 2.8% to 9.8%) annual increase in neonatal disease notification^7^.

There is evidence to suggest that the epidemiology of NTHi carriage and associated disease has been altered as a consequence of the introduction of pneumococcal conjugate vaccines (PCVs). This includes an increased NTHi incidence in children with AOM ^8-10^ as well as increased nasopharyngeal carriage in children who have received PCV ^11,12^ although it is important to note this is not a ubiquitous phenomenon^13^. In the UK two PCVs have been introduced in the routine childhood immunisation schedule, PCV7 (Prevenar 7™ introduced in 2006) and PCV13 (Prevenar 13™ introduced in 2010), that provide protection against particularly invasive capsular serotypes of *Streptococcus pneumoniae.* Importantly, they differ from PHiD10, a 10-valent pneumococcal conjugate vaccine used in some other countries, in that they are not conjugates of *Haemophilus* Protein D. Therefore, any effects on NTHi seen in the UK may be considered indirect rather than as a function of induced protective immunity to NTHi. Consequently, the underlying causes of these observations remain elusive. A limited understanding of the genomic-level population structure of NTHi has hindered efforts to elucidate impacts of PCV vaccination on carriage and disease. This would enable greater clarity regarding previous observations on NTHi co-occurrences with certain serotypes of *S. pneumoniae* ^14^. This paucity of data is of even greater importance given that the organism’s known high degree of genetic diversity, which is hampering the development of effective vaccines against NTHi^15^.

Here we use whole-genome sequencing of NTHi nasopharyngeal carriage isolates recovered from children <5 years of age recruited in this study across a five-year period and show that 1. The NTHi population exists in clearly delineated, temporally stable lineages, 2. That the introduction of PCV13 increased the carriage of NTHi, in the absence of apparent selection for distinct clones or lineages, and 3. Specific pneumococcal-NTHi lineage cocarriage associations exist that warrant further exploration.

## Results

### The introduction of PCV13 in the UK increased carriage of NTHi in children <5 years of age

Between the winters of 2008/09 and 2012/13 a total of 1569 nasopharyngeal swabs were taken from children <5 years of age attending outpatient clinics at a large UK hospital, University Hospital Southampton NHS Foundation Trust (NHS Research Ethics 06/Q1704/105). From these, 275 *Haemophilus* isolates were recovered (Table 1). Of these, 99.27% (n=273) were classified as NTHi using *in silico* capsular analysis. Three of these were *bexB/bexA* +/− and therefore classifiable as capsular deficient^16^. The two non-NTHi isolates were classified as serotype f.

**Table 1:** Carriage of Non-typeable *Haemophilus influenzae* and *Streptococcus pneumoniae* with age distribution of participants. NTHi carriage was significantly increased (*^1^ *p* < 0.05) in 2010/11 and 2012/12. No change in *S. pneumoniae* carriage was observed. The age of participants was significantly higher (***^2^ *p* < 0.001) in 2008/09 compared to all other years except for children who carried NTHi in 2011/12. Between children who carried NTHi age was significantly increased (***^3^ *p* < 0.001) in 2010/11 and 2011/12 compared to 2009/10. **4 The average age of NTHi carriers as significantly higher (*p* = 0.01) than that of all participants. §Age records for 37 participants were not available. ^a^ 95% CIs are shown in parentheses.

| Study Year | 2008/9 | 2009/10 | 2010/11 | 2011/12 | 2012/13 | Mean |
| --- | --- | --- | --- | --- | --- | --- |
| Participants (n) | 328 | 399 | 287 | 332 | 223 | 314 (310.8-317.2)<sup>a</sup> |
| **Bacterial Carriage** |  |  |  |  |  |  |
| NTHi (%) | 14.63 (n=48) | 14.79 (n=59) | 22.65 (n=65)<sup>*1</sup> | 16.57 (n=55) | 21.52 (n=48)<sup>*1</sup> | 18.03 (17.58-18.48)<sup>a</sup> |
| Spn (%) | 31.10 (n=102) | 27.82 (n=111) | 34.84 (n=100) | 29.73 (n=99) | 34.53 (n=77) | 31.6 (31.33-31.87)<sup>a</sup> |
| Co-colonised (NTHi + Spn) (%) | 6.71 (n=22) | 5.76 (n=23) | 8.71 (n=25) | 7.23 (n=24) | 10.76 (n=24) | 7.83 (7.48-8.18)<sup>a</sup> |
| **% Participants (n) by age<sup>§</sup>** |  |  |  |  |  |  |
| 0-2 | 0.61 (2) | 18.80 (75) | 7.32 (21) | 5.42 (18) | 2.24 (5) | 7.64 (24) |
| 3-12 | 25.61 (84) | 33.83 (135) | 34.15 (98) | 33.43 (111) | 45.29 (101) | 33.76 (106) |
| 13-24 | 26.52 (87) | 25.31 (101) | 25.78 (74) | 30.72 (102) | 31.39 (70) | 27.71 (87) |
| 25+ | 44.82 (147) | 21.05 (84) | 30.31 (87) | 28.92 (96) | 16.59 (37) | 28.66 (90) |
| **% NTHi Carriers (n) by age<sup>§</sup>** |  |  |  |  |  |  |
| 0-2 | 0.00 (0) | 6.78 (4) | 3.08 (2) | 1.82 (1) | 0.00 (0) | 01.82 (1) |
| 3-12 | 10.42 (5) | 32.20 (19) | 26.15 (17) | 20.00 (11) | 43.75 (21) | 27.27 (15) |
| 13-24 | 27.08 (13) | 27.12 (16) | 36.92 (24) | 30.91 (17) | 37.50 (18) | 32.73 (18) |
| 25+ | 60.42 (29) | 33.90 (20) | 33.85 (22) | 49.09 (27) | 16.67 (8) | 38.18 (21) |
| **Avg. Age All Participants (months)** | 25<sup>***2</sup> | 15 | 20 | 20 | 16 | 19<sup>**4</sup> (18.80-19.20)<sup>a</sup> |
| **Avg. Age NTHi Carriers (months)** | 31<sup>***2</sup> | 19 | 22<sup>***3</sup> | 26<sup>***3</sup> | 18 | 23<sup>**4</sup> (22.34-23.633)<sup>a</sup> |

Carriage prevalence for NTHi ranged from 14.63% in 2008/9 to 22.65% in 2010/11 with a mean of 18.03% (95% CI 17.58 – 18.48) (Table 1 and Figure 1). Using a multivariable, binomial logistic regression model there was higher carriage prevalence of NTHi in the post-PCV13 years of 2010/11 and 2012/13 (*p* < 0.05), but not 2011/12, compared to the pre-PCV13 years. Children that carried NTHi were older (*p* < 0.05) (Table 1 and Supplementary Figure 1) and every increase in age by one month was significantly associated with increased odds of carriage (OR 1.02, *p* < 0.001). Examining each year, in 2008/09, the first year of the study presented here, there was a significantly older population recruited compared to all other years (ANOVA with Tukeys HSD *p* < 0.001, Table 1).

**Figure 1:**
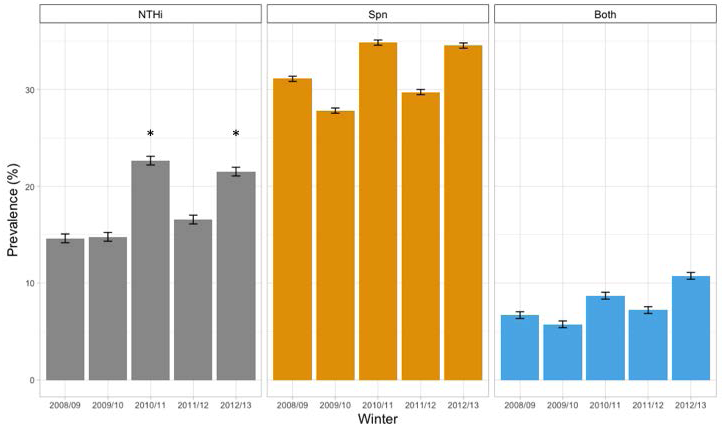
Carriage Prevalence of Non-typeable *Haemophilus influenzae* and *Streptococcus pneumoniae*. NTHi carriage (grey bars) ranged from 14.63% in 2008/9 to 22.65% in 2010/11 (average of 18.03%). Using a binomial logistic regression model, significantly higher carriage prevalence of NTHi in 2010/11 and 2012/13 (* *p* < 0.05) was observed. Neither *S. pneumoniae* carriage (orange bars) nor co-carriage (blue bars) increased significantly in this period. Error bars – 95% confidence interval.

Co-carriage of pneumococci and NTHi varied from 5.76% in 2009/10 to 10.76% in 2012/13, although this was not significantly different between years. Presence of S. *pneumoniae* however was significantly associated with NTHi carriage (*p* < 0.001, OR 1.92, 95% CI 1.66-2.17). Comparing NTHi co-carriage with *S. pneumoniae* vaccine serotype (VT) versus non-VT in 2008/09 and 2009/10 (the years preceding PCV13 introduction) revealed significantly increased odds of co-carrying VT pneumococci (*p* < 0.05, OR 2.36, 95% CI 1.17-4.75). The three most common co-carried VT serotypes were 19A (n=19), 6A (n=11) and 6B (n=6).

In total 119 NTHi individual multi-locus sequence types (MLSTs) were identified (Figure 2). MLST diversity, as measured using Simpsons (1-D), within each year was high, ranging from 0.97 in 2008/09 to 0.99 in 2010/11 and 2011/12 respectively. To determine whether there had been an increase in NTHi MLST diversity following PCV13 introduction the first two and the last two years were grouped as pre- and post-PCV13 eras respectively. The middle year of 2010/11 was excluded in this analysis as we hypothesised this may represent an interim perturbed NTHi population not truly reflective of a post-PCV13 era. Although a slight increase in diversity was noted (Figure 2) this was not statistically significant.

**Figure 2:**
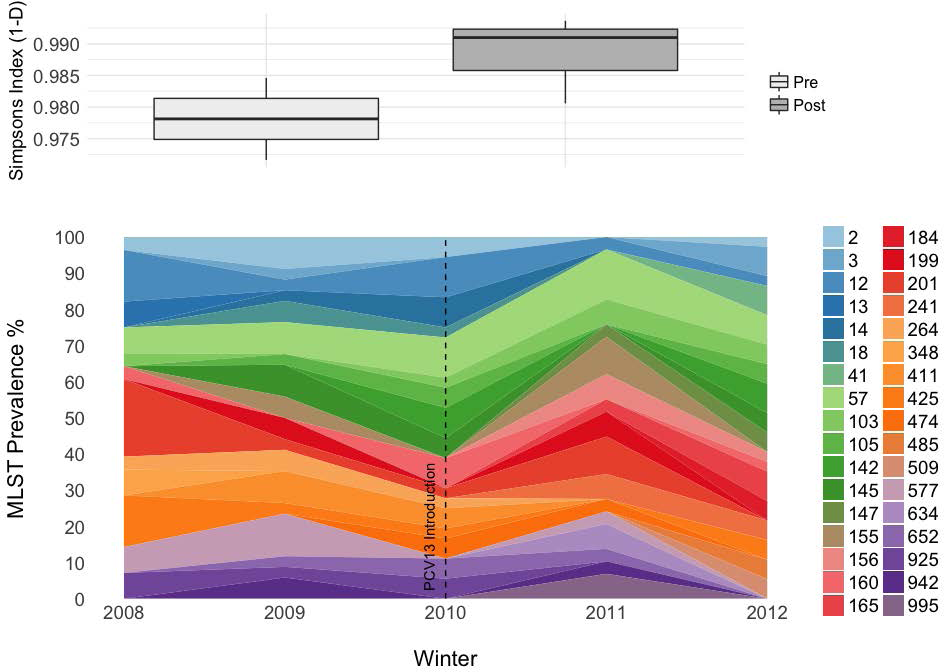
NTHi MLST Prevalence and Diversity During PCV Implementation. For visual clarity MLST percentage prevalence (A.) is only shown for sequence types that were identified at least twice in any given year of the study. The dashed line delineates the pre- and post-PCV13 eras that were used to calculate MLST diversity (as measured by Simpsons 1-D) and shown in B. Error bars – 95% confidence intervals.

### The NTHi population can be defined by 11 discrete and temporally stable lineages

Figure 3 shows the population structure for 266 NTHi carriage isolates for which genomic data was available (those excluded having a sequencing depth equating to less than 30-fold coverage) in combination with an additional 89 isolates which had previously been used to identify six phylogenetic clusters^17^. The total core genome alignment length represented only 313 Kbp (16%) of the on average 1.92 Mbp genome. HierBAPS analysis revealed eleven lineages of NTHi. The polyphyletic lineage 9 was the most predominant (n=54; 19.6% of the carriage isolates) with the fewest being five isolates belonging to lineage 2. All of the 89 additional isolates were classified concordantly with their six previously identified clusters, but with additional delineation. Isolates belonging to cluster IV were further separated into lineages 2 and 11, cluster V into lineages 8 and 4, and cluster VI as lineages 6,7 or 9 (Supplementary Data 1).

**Figure 3:**
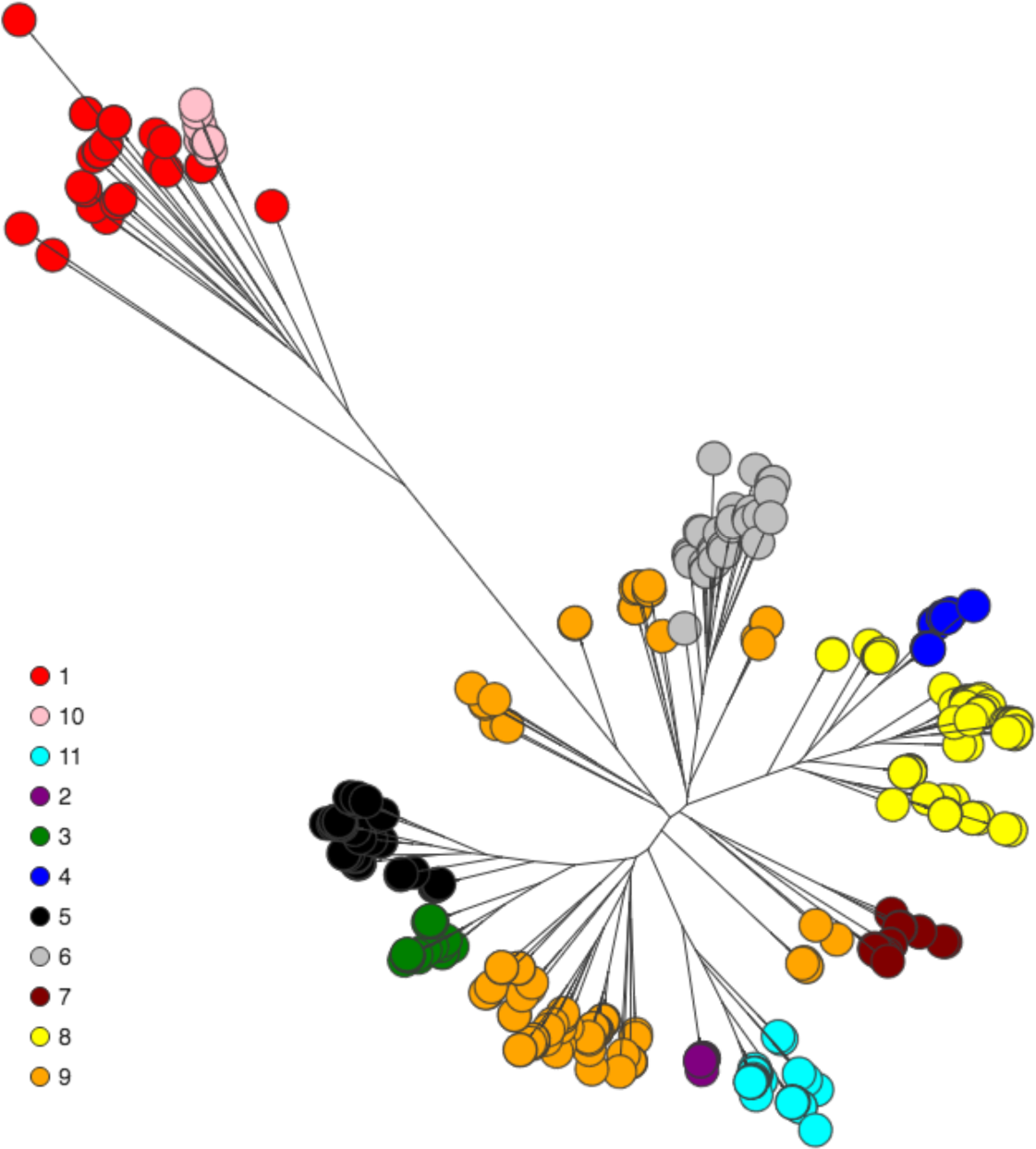
Population structure of the NTHi carriage isolates as determined by hierBAPS analysis. Terminal nodes of the maximum-likelihood core genome tree are coloured according the eleven lineages identified.

We sought to address the paradox of an apparently highly diverse collection of NTHi, based on number of STs identified, within relatively few lineages. We hypothesised that the intra-lineage diversity is balanced by a stable population structure at a higher level of clustering. Here Bray-Curtis dissimilarity was used to account for both lineage presence and abundance between years. As shown in Figure 4 (and Supplementary Figure 2), lineages exhibited very little difference between years. Any variation was not due to PCV13 introduction as the most dissimilar years were both post-PCV13 introduction, 2010/11 and 2012/13 (0.41 Bray-Curtis dissimilarity index).

**Figure 4:**
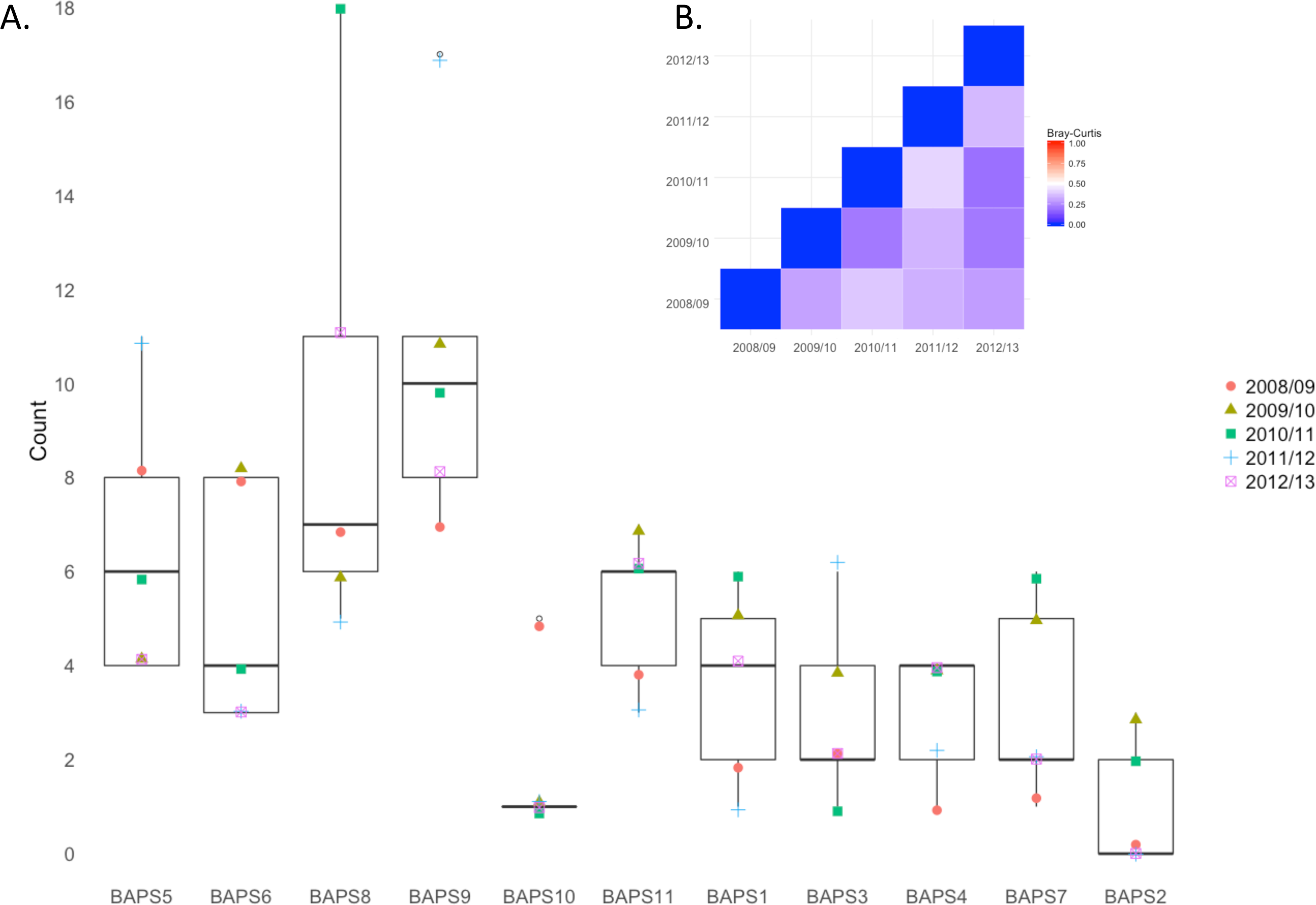
A. Box and whisker plot showing NTHi hierBAPs lineage occurrence across years. Lineages are ranked according to the abundance in 2008/09. Coloured symbols represent each of the five sampling periods 2008/09 to 20012/13. B. Heat-map showing Bray-Curtis dissimilarity index based on presence and abundance of hierBAPS lineages within each year of the study. Red indicates less similar populations with blue showing highly similar structuring.

MLST diversity within lineages varied considerably (Figure 5) ranging from 30 unique MLSTs from 54 lineage 9 isolates, which was also the most frequently observed, to lineage 2, which consisted of just five isolates of ST411. The most common STs encountered were ST57 (n=15, lineage 11), ST201 (n=11, lineage 5) and ST12 (n=11, lineage 8). Lineage prevalence was not a good predictor of intra-lineage diversity measured by Simpsons 1-D (Table 2) r(10) = 0.122, *p* > 0.5.

**Figure 5:**
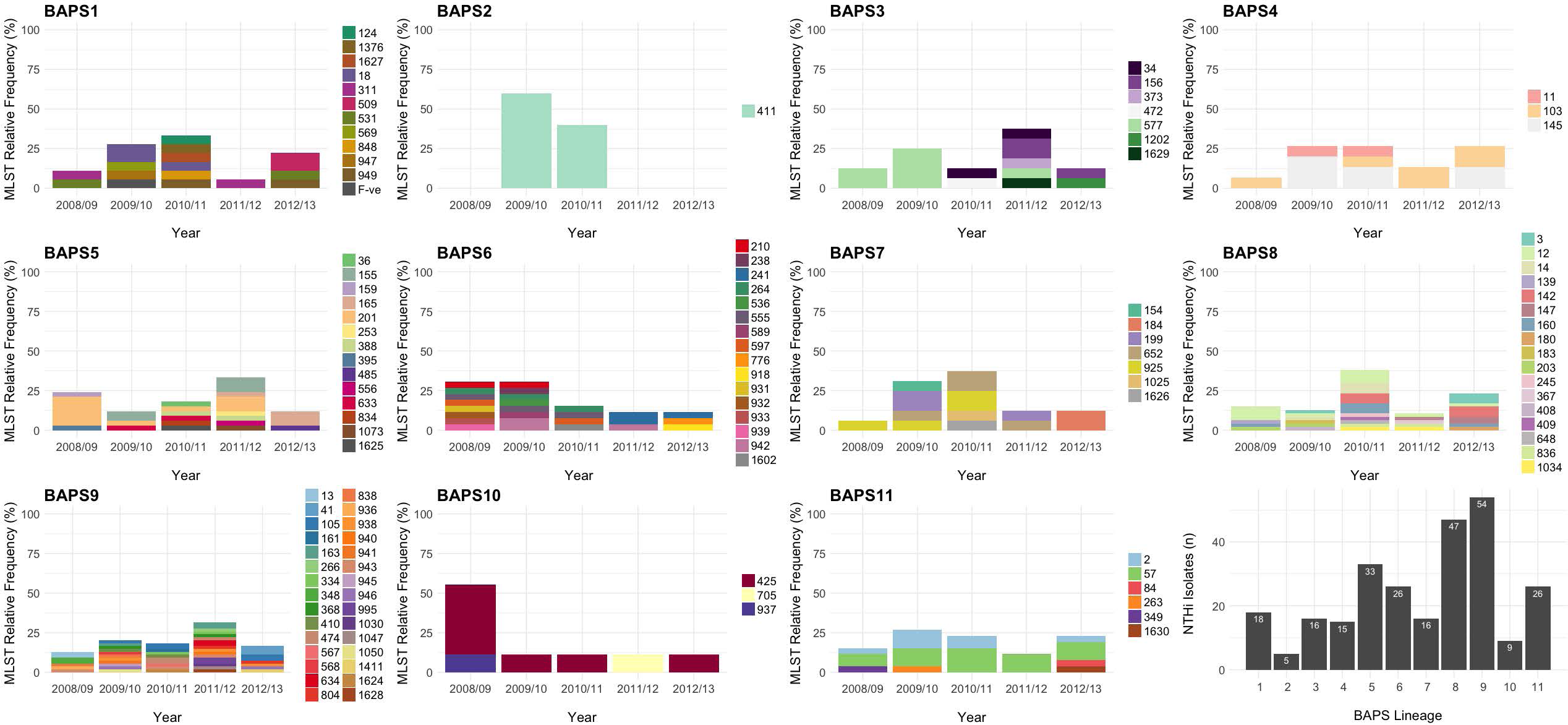
Intra-lineage diversity of MLSTs in NTHi. The distribution of each MLST uniquely associated with a hierBAPS lineage (BAPS1-11) is shown. The cumulative isolate number for each lineage for all five years is also given in the bottom right plot.

**Table 2:**
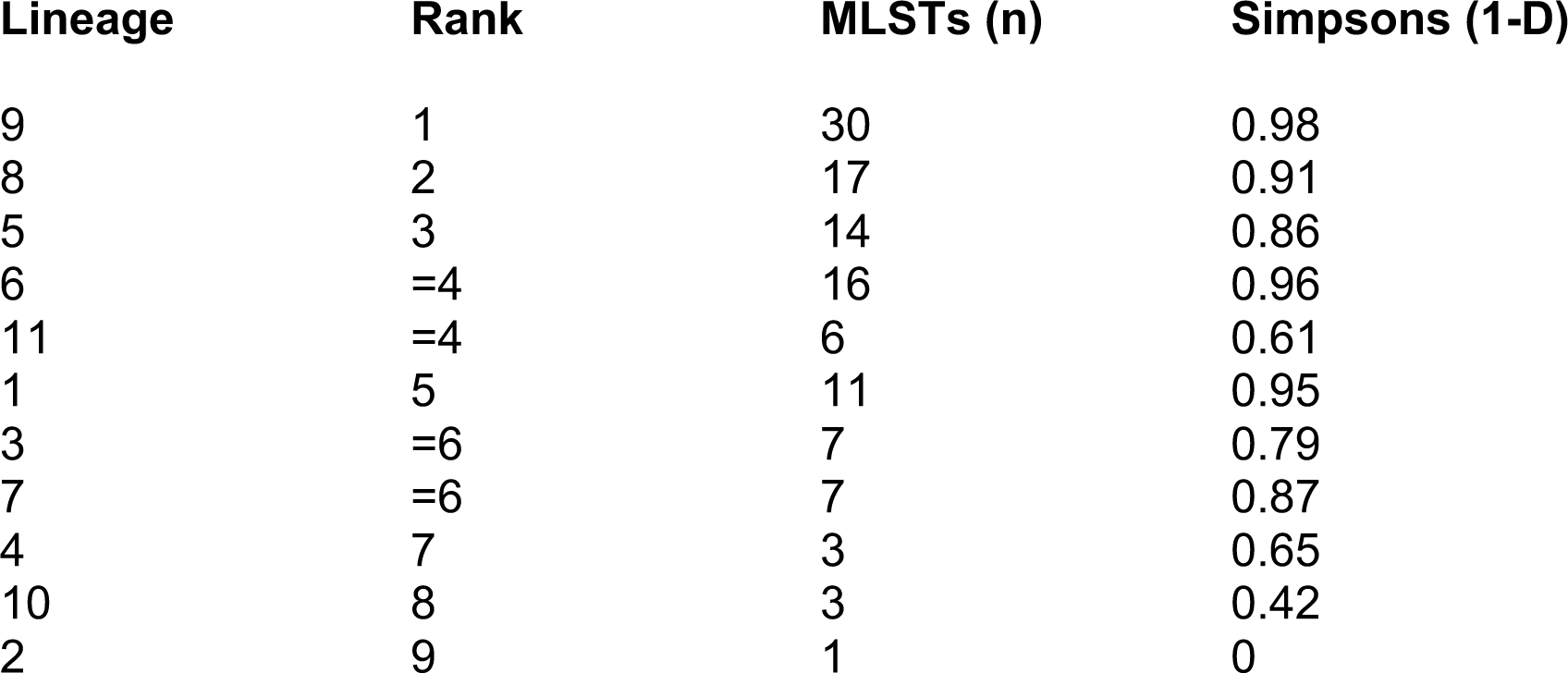
MLST Diversity within each hierBAPS lineage. Rank is based on cumulative abundance from years 2008/09 to 2012/13.

### NTHi lineage 6 was significantly associated with carriage of *S. pneumoniae* pre-PCV13

In the years preceding and following PCV13 introduction there was little change in the co-carriage of any one particular NTHi lineage with *S. pneumoniae* (Figure 6). However, lineage 6 had an OR for co-carriage of 14.75 (95% CI: 3.14-69.38) and 16.95 (95% CI: 0.93-309.96) in these two periods, with the former being statistically significant (*p* < 0.001) indicating a correlation between the carriage of this lineage of NTHi and *S. pneumoniae;* co-carriage was found for 87.5% and 100% isolates respectively Interestingly these increased odds were significant in the two years prior to PCV13 until 2010/11, when we predict the impact of PCV13 may have begun, whereupon this relationship was altered (Figure 6). Pre-PCV13, 31.3% (5/18) of lineage 6 isolates were co-carried with vaccine serotype pneumococci (serotypes 6B, 19A and 7F). In comparison only 8, lineage 6 isolates were found in the following three post-PCV13 years. The absence of VT pneumococci in this period coupled with a lower isolate number leaves this potential impact of PCV13 open to interpretation. By contrast, in the pre-PCV13 era lineage 11 exhibited a negative association with *S. pneumoniae* (OR 0.12 95% CI: 0.05-1.01 *p* = 0.05) however this was not statistically significant in the post-PCV13 era.

**Figure 6:**
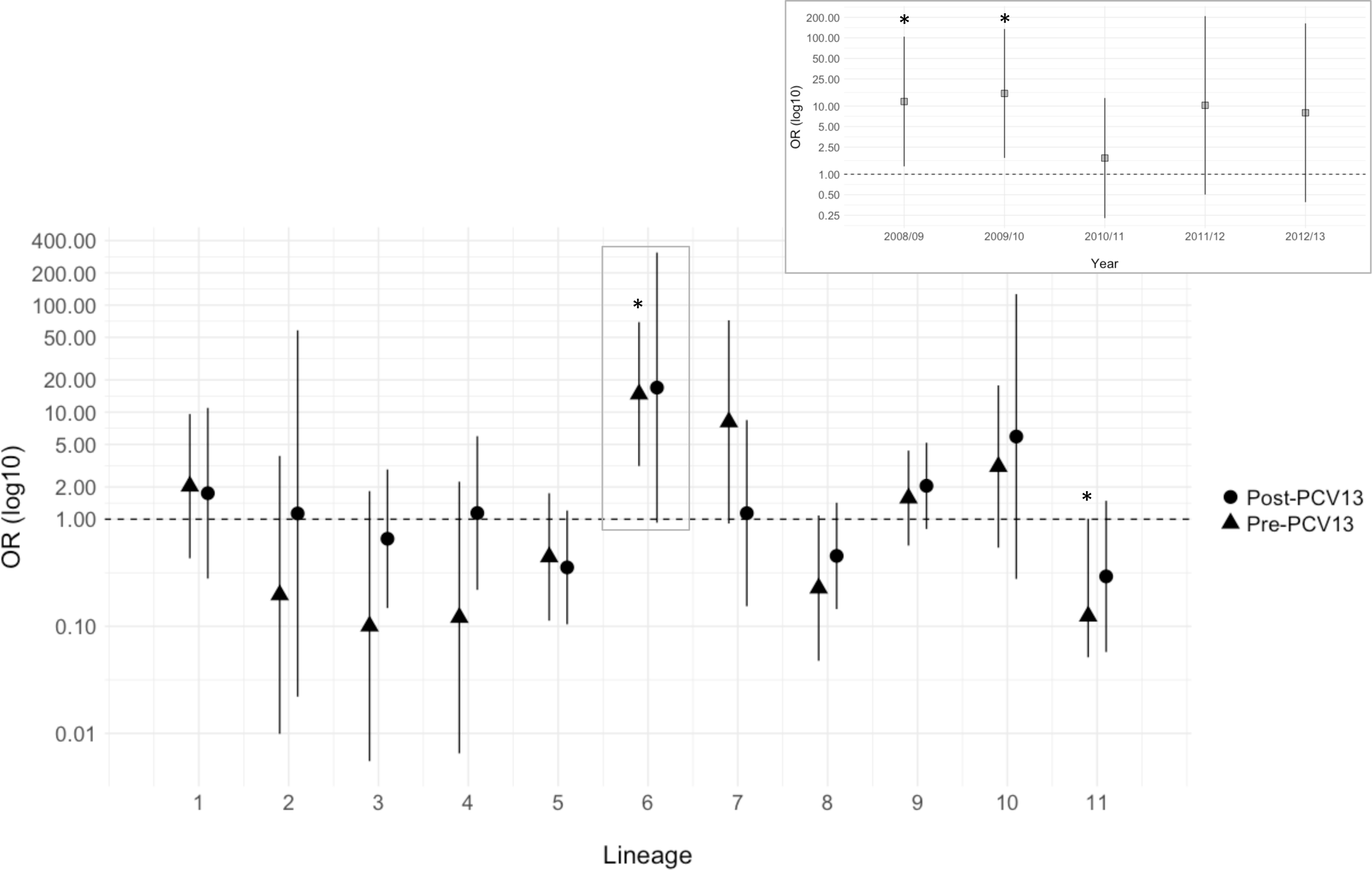
Odds of co-carriage of *S. pneumoniae* for each NTHi lineage pre- and post-introduction of PCV13. Here odds ratios were calculated using data from 2008/09-2009/10 as pre-PCV13 and 2011/12-2012/13 as post-PCV13. Lineage 6 pre-PCV13 was significantly (* *p* = 0.007) associated with carriage of *S. pneumoniae.* In the same period lineage 11 was significantly (* *p* = 0.05) associated with non-carriage of *S. pneumoniae.* These two associations were not present in the post-PCV13 era. Inset is shown the odds ratio for co-carriage of *S. pneumoniae* for lineage 6 across all years.

### Diverse levels of recombination exist between NTHi lineages

Within the lineages identified here an almost universally high degree of recombination was observed (Table 3). Although recombination occurred on average 25% less often than mutation (P/θ 0.742, 95% CI: 0.713-0.771), it produced 19 times (r/m 19.075 95% CI: 17.38-20.77) more substitutions than *de novo* mutation. Exceptions included lineages 1, 7 and 9 which had significantly lower relative impacts of recombination with r/m values of 3.467 (*p* = 0.0002), 10.642 (*p* = 0.0267) and 4.651 (*p* = 0.0005) respectively. In addition, lineage 10 had a P/θ of 1.103 indicating recombination occurred approximately 10% more often than mutation and this caused 56 times more substitutions. MLST diversity within each lineage was correlated to r/m such that lineages with lower Simpsons 1-D (and thus greater diversity) were shown to also have higher r/m (*r*(10) 0.733 *p* < 0.001).

**Table 3:** Extent of recombination identified in the eleven NTHi lineages. Output from ClonalFrame ML is shown. R/θ = ratio of recombination to mutation, δ = the mean length of recombination imports, ν = average divergence of imports, r/m = the relative effect of recombination to mutation. Higher r/m values indicate the change in rate at which a nucleotide is likely to be altered due to recombination compared to de novo mutation. Lineages 1 and 9 had significantly lower r/m compared to the population mean whereas lineage 10 was significantly higher (p values based on z test of population means).

| Lineage | P/θ | δ (bp) | v | r/m | p (r/m) |
| --- | --- | --- | --- | --- | --- |
| 1 | 0.854 | 88 | 0.046 | 3.47 | 0.0002* |
| 2 | 0.184 | 3226 | 0.020 | 11.93 | 0.0521 |
| 3 | 0.831 | 892 | 0.031 | 23.11 | 0.8204 |
| 4 | 0.550 | 1343 | 0.039 | 28.74 | 0.9860 |
| 5 | 0.848 | 782 | 0.030 | 19.56 | 0.5440 |
| 6 | 0.904 | 641 | 0.027 | 15.51 | 0.2088 |
| 7 | 0.727 | 495 | 0.030 | 10.64 | 0.0276* |
| 8 | 0.809 | 730 | 0.031 | 18.31 | 0.4312 |
| 9 | 0.534 | 277 | 0.031 | 4.65 | 0.0005* |
| 10 | 1.103 | 1449 | 0.036 | 56.83 | <0.0001* |
| 11 | 0.819 | 692 | 0.030 | 17.07 | 0.3240 |
| Mean | 0.742 | 965 | 0.031 | 19.075 | - |
| 95% CI | 0.771-0.713 | 865-1064 | 0.031-0.032 | 17.38-20.77 | - |

### Recombination hot-spots are characterised by involvement in metabolic / biosynthetic pathways

Recombination across the core genome is shown in Figure 7A and is mapped against a maximum-likelihood core genome phylogeny. Here recombination is shown by dark blue and mutation as white. The variability between lineages is clear, as is the non-uniformity of recombination across the core genome alignment. This variability is illustrated in Figure 7B where recombination blocks are counted and plotted according to position in the alignment. Each of the eleven lineages were also analysed separately, these can be found as Supplementary Figures 3 to 13.

**Figure 7:**
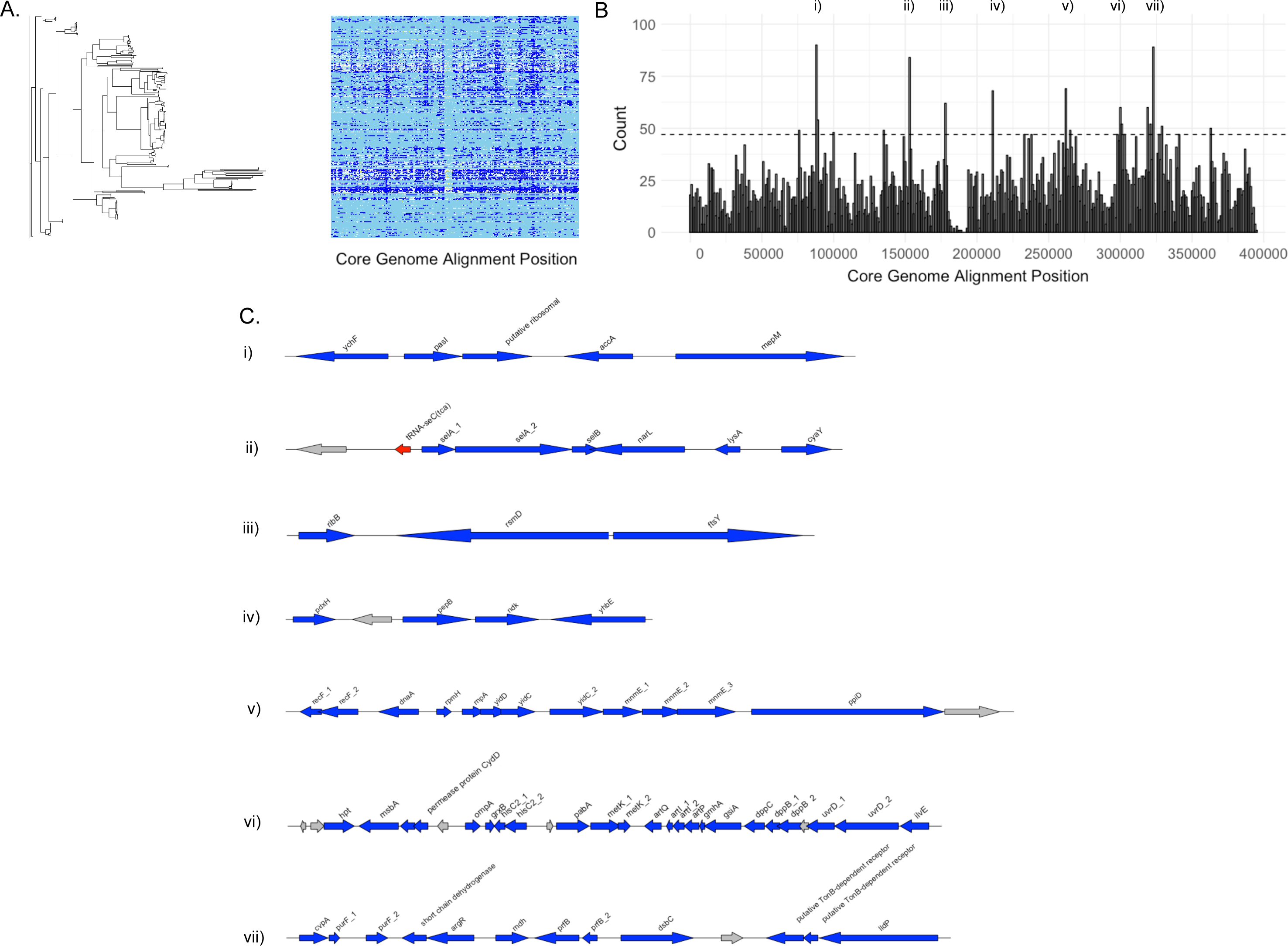
Recombination in the core genome of NTHi. Maximum-likelihood phylogeny and a recombination plot using ClonalFrameML (A) is shown adjacent to a density plot of recombination blocks across the core genome alignment (B). Here seven regions that had a density of recombination blocks above the 95^th^ quantile are shown. Annotations are shown in C, with grey arrows denoting hypothetical proteins, red - tRNAs and blue - identifiable coding genes.

Seven regions are shown to contain recombination counts above the 95% inter-quartile range. Annotations of these are shown in Figure 7C. The predominant characteristic of these loci is the significant representation of genes involved in metabolic and biosynthesis pathways. These include selenoprotein biosynthesis (*selA1-2, selB*), amino acid metabolism and scavenging (*pepB, ilvE, hisC2, grxB, art, hpt, cydD*), carbon source utilisation (*metK1/2, lidP*) and protein formation and transport *(dsbC, yhbE, msbA, yidD/C, ppiD).* Several genes that a play role in responses to nutrient availability / stress were also noted and included *narL* which mediates nitrate-response transcriptional regulation^18^, the DNA repair-associated genes *recF1* and *recF2,* and adenylate cyclase (*cyaD*) which controls competence. Of note is that only one gene associated with outer membrane proteins was identified within these seven regions; *ompA* which encodes outer membrane protein P5.

### Prevalence of antibiotic resistance genes is low except for β-lactamase-negative ampicillin resistance (BLNAR) mutations in ST411 of lineage 2

The identification of antibiotic resistance genes is shown in Figure 8. Alleles for spectinomycin resistance and the multi-drug efflux pump, *hmrM* were found to be near ubiquitous at 96% (n=257) and 98% (n=262) of isolates respectively. APH(3) alleles for aminoglycoside resistance were found in less than 2% (n=5) of isolates which included STs from lineages six (ST932 and 264) seven (ST154), eight (ST3 and 142) and nine (ST1411). One of these isolates (a lineage eight, ST3 from 2012/13) was also positive for *catII* (chloramphenicol), *tet* and *tetD.* The same isolate carried resistance to sulphonamide *(sul2)* as did one other, a lineage seven, ST154 from 2008/09.

**Figure 8:**
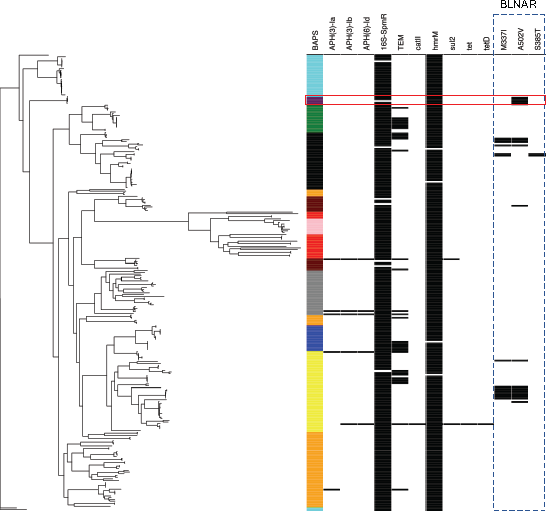
Identification of antimicrobial resistance genes and mutations in NTHi. The maximum-likelihood phylogeny (left) is annotated by coloured block according to hierBAPS lineage designation (column 1). Presence of loci / mutations associated with resistance are shown in black. The mutations in the Penicillin Binding Protein 3 (PB3) encoding *ftsl,* and associated with high level β-lactamase-negative ampicillin resistance, are highlighted by the dashed rectangle. The red box shows lineage 2 as being the only lineage in which all isolates harboured a mutation associated with this phenotype.

Of the 34 isolates in which β-lactamase resistance (TEM) was identified, seven belonged to lineage 3 (ST577 n=6, ST156 n=1) six of lineage 4 (ST103), five from lineage five (ST165 n= 4) and two from each of lineages six, seven and nine. The majority however were from lineage 8, four of which were ST160.

β-lactamase-negative ampicillin-resistance (BLNAR), a consequence of mutations in *ftsl* that encodes the transmembrane component of penicillin binding protein (PBP) 3, was found in approximately 35 isolates and limited to four lineages (Figure 8). The mutations that have previously been characterised as producing high-level, group III, resistance^19^, and identified here, included the Methionine to Isoleucine (M337I) in 5.6% of isolates (n=15), Alanine to Valine (A502V) in 7.5% (n=20), and Serine to Threonine (S385T) in just two isolates (ST155, lineage 5). None harboured the Leucine to Phenylalanine conversion at position 389. The majority of the former was identified in multiple STs of lineages eight and five. Interestingly all lineage two isolates (ST411, n= 5) harboured the A502V mutation.

## Discussion

Non-typeable strains of *Haemophilus* are recognised to cause significant human disease including community acquired pneumonia ^20^. We explored epidemiological shifts in NTHi carriage over a five-year period that included PCV13 replacement of PCV7 in the UK’s National Immunisation Programme in 2010. We demonstrate increased carriage of NTHi in children <5 years of age following this time point, which was not linked to the expansion of any one particular clone. In addition, to our knowledge this is the first study to demonstrate the temporal stability of NTHi lineages pre- and post the introduction of a PCV. We have shown the existence of a discrete lineage structure that is stable over time and which did not measurably shift in response to PCV13. The clonality of the population was concordant with previous examinations of population structure based on MLST^21^ and whole genomes of geographically diverse isolates collections^17^. The hierarchical analysis we used gave greater phylogenetic clarity and revealed hitherto unrecognized resolution of the lineages of NTHi.

We also observed that in the pre-PCV13 era, NTHi carriage was more associated with carried pneumococcal serotypes that would ultimately be targeted by PCV13 (19A, 6 and 11) compared to non-VT strains. Data from previous studies have shown similar trends demonstrating that PCV13 serotypes were more associated with *Haemophilus* carriage in contrast to non-PCV13 serotypes^14^. This suggests there are serotype-dependent carriage dynamics between these bacterial species. A plausible outcome of vaccination is that PCV disrupts these associations. We believe this is the first description of specific NTHi lineages that exhibit notable interactions with pneumococci, regardless of whether these are VT serotypes. The fact that there could be NTHi strains that differ in the direction and amplitude of interaction with pneumococci is not surprising as there are many examples of competition, antagonism and synergism between the two genera ^22-25^ suggestive of a complex relationship. Specifically, co-carriage likelihoods per lineage were not consistent for *S. pneumoniae,* whereby lineage 2 was associated with increased likelihood of pneumococcal carriage and the converse true for lineage 11. However, we were unable to determine the NTHi genetic basis of the observed relationships between lineages 6 and 11 with pneumococci. This was due to a low number of isolates; lineage 6 for example represented less than 10% of the NTHi isolates collected in this study. These low numbers of isolates render the application of genome-wide association studies virtually impossible. In addition the phenotypes are not clearly delineating (for example as is antibiotic resistance) and they reside within a monophyletic lineage that pose a particular challenge to association studies of this nature (although methods to account for this within bacterial populations do exist)^26^.

Recombination varied considerably in this population and correlated to intra-lineage diversity. Recombination has previously been shown to be a significant driver of *Haemophilus* sp. genetic diversity ^17,21^, modulation of outer-membrane proteins ^27^, and transfer of antibiotic resistant genes ^28^. The high levels of recombination observed here, both in terms of frequency and locus involvement are in keeping with that observed by Cody *et al.,* (2003) who found recombination in genes associated with LPS biosynthesis and as well as housekeeping.

There are a number of limitations to this study. Firstly, the sampling is limited to one geographical region and a defined, narrow subset of the population (children <5 years of age). The extrapolation of the genomic population structure detailed in this study to wider isolate collections must therefore be done with caution, although similar, low numbers of lineages have previously been identified from much broader isolate collections ^17,21,29^. The most striking feature of temporal stability is one that can be addressed in future with continued sampling from our paediatric outpatient population. Lastly these isolates are all from carriage and may not be representative of the genotypic diversity associated with disease. It will be interesting for future work to compare the diversity between strains isolated from chronic and acute pathologies from both children and older age cohorts.

The use of PCVs have radically reduced the burden of invasive pneumococcal disease (IPD) both in the UK^30^ and globally ^31-37^. In the UK, this IPD reduction has occurred in the absence of any loss of overall pneumococcal carriage prevalence ^38,39^, and is a consequence of serotype replacement. The lack of penetrance between carriage replacement with non-VT serotypes and IPD is a consequence of lower invasiveness in the non-VT pneumococci ^40^. Regardless, the replacement of VT pneumococci represents a disruption in nasopharyngeal microflora. Evidence for indirect effects associated with PCV on *Haemophilus* disease and carriage have been noted ^8-12^ and PCV vaccination in the very young (<12 months) has been shown to cause a more disordered nasopharyngeal microbiota^41,42^. We hypothesise that this niche disruption, in combination with the adaptation of *S. pneumoniae* to maintain its relative fitness in the face of selective pressures in response to PCV introduction (Red Queen dynamics), ^*43*^ impacted the interactions between pneumococci and other microbiota resident in the nasopharynx. The disruption appears to manifest as the general, non-specific flux in carriage prevalence of NTHi (as the dominant *Haemophilus* sp.) seen here. The ramifications of there being particular lineages of NTHi that have extreme competitive or commensal relationships with pneumococci are important clinically in populations. Specifically, the impact of PCV on nasopharyngeal microbiomes will influence future conjugate vaccine design and use as well as interpretation of experimentally derived interaction models.

This study represents the largest NTHi genomics study to date. Through this analysis we have shown that the introduction of PCV13 directly influenced the epidemiology of NTHi by increasing carriage prevalence in a young paediatric population. This included altered associations between specific genomic lineages of NTHi with pneumococci. Although highly diverse at the MLST level, the eleven lineages identified through genome analysis displayed remarkable temporal stability during this period. Our findings have clinically important implications for the implementation of future pneumococcal vaccinations as well as the design of any future *Haemophilus* vaccine candidates.

## On-line Methods

### Bacterial Isolates

Nasopharyngeal swabs were collected from children aged <5 years of age who attended outpatient clinics at Southampton General Hospital during five consecutive winters, October to March, 2008/9 to 2012/13. The Southampton and South West Hampshire Research Ethics Committee, B approved this study (NHS Research Ethics 06/Q1704/105). This study, designed to characterize *S. pneumoniae* carriage, has been described in detail elsewhere^38,39,44^. Informed consent was obtained before or after an outpatient appointment, no specific outpatient clinic was targeted. The participant was either the child attending clinic or their sibling. Only one child per family was swabbed. Age was the primary exclusion criteria. Rayon tipped Transwabs (Medical Wire, Corsham, UK) in charcoal Amies media were plated onto Chocolate agar with Bacitracin within 9 hours of collection at the Health Protection Agency Southampton Laboratory (now part of Public Health England) between 2008/09 and 2011/12 and by technical staff in our research group during 2012/13. Presumptive *Haemophilus* were plated onto blood agar with X (hemin), V (nicotinamide adenine dinucleotide, NAD) and XV discs (Oxoid, UK). Presumptive *Haemophilus influenzae* were determined by growth around XV discs only. Only one colony of *Haemophilus* per participant swab was selected for further analysis.

### DNA Extraction and Quantification

Genomic DNA was extracted from a sweep of NTHi colonies using a QIAmp DNA Mini kit (Qiagen, UK) as per manufacturer’s instructions. The concentration of genomic DNA was determined using Qubit™ 2.0 fluorometric quantitation (Thermo-Fisher, UK).

### Whole Genome Sequencing

Sequencing was done using Illumina MiSeq, with V2 chemistry to generate 2 x 250 bp paired-end read data to a depth of approximately 30-fold coverage for each bacterial isolate.

### Genome Assembly and Annotation

Paired-end reads were trimmed using trimmomatic v0.32^45^ and *de novo* assembled using SPAdes v3.10.1^46^. Contiguous sequence orientation and gap filling to create scaffolds was undertaken using the assembly_improvement script from the Wellcome Trust Sanger Institute (http://github.com/sanger-pathogens/assemblyimprovement). Annotation was done using Prokka v1.10^47^.

### MLST and Capsular Loci Identification

Read mapping for MLST designation was done using SRST2 v0.1.5^48^. Confirmation of capsular status was done by *in silico* PCR in exonerate v2.2.0 (http://github.com/nathanweeks/exonerate) using previously published primers^16^.

### Antibiotic Resistance Gene Identification

Identification of antibiotic resistance determinants was done using Ariba^49^ v2.10.1. using the Comprehensive Antibiotic Resistance Database ^50^. Identification of β-lactamase-negative ampicillin-resistance (BLNAR) due to mutations in *ftsl* was done using *in silico* primers SSNF2 and KTGR2 as previously described^19^. Protein alignments were done in Seaview v4.5.4. Results were visualised using Phandango (http://phandango.net/)^51^.

### *Haemophilus influenzae* Genomes

Genomes as used by De Chiara et al. (2014)^17^ were downloaded from ftp://ftp.sanger.ac.uk/pub/project/pathogens/Haemophilus/influenzae/NTstrains/

### Analysis of Population Structure

Core genome alignments were constructed using parsnp from harvest v1.2^52^ with -x to filter recombination. Here the minimum MUM anchor was set to 17 to force alignment of more dissimilar genomes. The resultant xmfa was converted to fasta using the script xmfa2fasta.pl (http://github.com/kjolley/seqscripts/blob/master/xmfa2fasta.pl) and Gblocks v0.91b^53^ was used to remove contiguous non-conserved regions from the alignment. A maximum likelihood phylogeny was constructed using the CIPRES hosted RAxML-HPC v8 ^54,55^ with a General Time Reversible model, GAMMA substitution rate and rapid bootstrapping (n=100). Population structure was determined using hierBAPS^56^ implemented with four hierarchy levels and an upper cluster limit of 20. The resultant tree and metadata was visualised using Microreact (http://microreact.org/) ^57^.

### Recombination

Recombination was determined using ClonalFrameML^58^ v1.11 with -emsim set at 100 simulations.

### Statistical Analyses

All statistical analyses were done in R Studio v3.4.0. Generalised linear models were done using the function *glm()* with ‘family=binomial’, linear models were done using function *lm*().

### Genomic Data

Data is deposited in the European Nucleotide Archive (ENA) with accession numbers: TBC

## Acknowledgments

We wish to thank staff at the Southampton NIHR Wellcome Trust Clinical Research Facility for their contribution towards the collection of samples and Public Health England microbiologists for microbiological processing of swabs. We gratefully acknowledge both the guardians and participants that made this study possible. This study was made possible via investigator-initiated research grants from Pfizer to SCC and SNF for which we also acknowledge the contributions in initial research development from Dr Johanna M. Jefferies.

## Contributions

SCC and SNF conceived and secured funding for the study. SCC and SNF were the site primary investigators. DEM, VTD and KLO undertook participant recruitment and microbiological isolation. VTD carried out DNA extractions, sequencing and genome assembly. DWC analysed the whole genome sequencing data and wrote the manuscript. All authors reviewed the manuscript.

---

Supplementary Figure 1: Histogram of participant’s age (left) and of NTHi carriers (right).

Supplementary Figure 2: Prevalence of each NTHi hierBAPS lineage across study years.

Supplementary Figure 3: Recombination in the core genome of hierBAPS lineage 1. Maximum-likelihood phylogeny and a recombination plot using ClonalFrameML is shown above a density plot of recombination blocks across the core genome alignment.

Supplementary Figure 4: Recombination in the core genome of hierBAPS lineage 2. Maximum-likelihood phylogeny and a recombination plot using ClonalFrameML is shown above a density plot of recombination blocks across the core genome alignment.

Supplementary Figure 5: Recombination in the core genome of hierBAPS lineage 3. Maximum-likelihood phylogeny and a recombination plot using ClonalFrameML is shown above a density plot of recombination blocks across the core genome alignment.

Supplementary Figure 6: Recombination in the core genome of hierBAPS lineage 4. Maximum-likelihood phylogeny and a recombination plot using ClonalFrameML is shown above a density plot of recombination blocks across the core genome alignment.

Supplementary Figure 7: Recombination in the core genome of hierBAPS lineage 5. Maximum-likelihood phylogeny and a recombination plot using ClonalFrameML is shown above a density plot of recombination blocks across the core genome alignment.

Supplementary Figure 8: Recombination in the core genome of hierBAPS lineage 6. Maximum-likelihood phylogeny and a recombination plot using ClonalFrameML is shown above a density plot of recombination blocks across the core genome alignment.

Supplementary Figure 9: Recombination in the core genome of hierBAPS lineage 7. Maximum-likelihood phylogeny and a recombination plot using ClonalFrameML is shown above a density plot of recombination blocks across the core genome alignment.

Supplementary Figure 10: Recombination in the core genome of hierBAPS lineage 8. Maximum-likelihood phylogeny and a recombination plot using ClonalFrameML is shown above a density plot of recombination blocks across the core genome alignment.

Supplementary Figure 11: Recombination in the core genome of hierBAPS lineage 9. Maximum-likelihood phylogeny and a recombination plot using ClonalFrameML is shown above a density plot of recombination blocks across the core genome alignment.

Supplementary Figure 12: Recombination in the core genome of hierBAPS lineage 10. Maximum-likelihood phylogeny and a recombination plot using ClonalFrameML is shown above a density plot of recombination blocks across the core genome alignment.

Supplementary Figure 13: Recombination in the core genome of hierBAPS lineage 11. Maximum-likelihood phylogeny and a recombination plot using ClonalFrameML is shown above a density plot of recombination blocks across the core genome alignment.

